# Hippocampal time cell dynamics evolve with learning to reflect cognitive demands

**DOI:** 10.1101/2025.05.20.655191

**Authors:** Erin R. Bigus, John C. Bowler, Dua Azhar, Dylan Zukowsky, James G. Heys

## Abstract

The hippocampus creates cognitive maps, or internal representations that reflect knowledge of the external world. Hippocampal time cells are thought to represent the temporal structure of experiences, or temporal context. However, it remains unknown whether hippocampal time cells display learning dynamics that reflect increased knowledge of the temporal relationships that define a context. To address this gap, we utilized a behavioral paradigm with a shaping curriculum that allows animals to systematically acquire knowledge of temporal structure, ultimately enabling them to perform a temporal Delayed Non-Match to Sample (tDNMS) task. We conducted two-photon calcium imaging on large populations of CA1 neurons as mice progressed through the curriculum—from their initial exposure to the task structure to their successful discrimination of context at the end of training. Time cells were present from the outset, yet their activity evolved with experience, both at the single-cell level and across the population. Notably, at key moments in the curriculum, time cell dynamics adapted to reflect whether mice generalized across contexts or discriminated between them. Our findings suggest that CA1 time cells not only represent temporal context but may also reflect the processes by which temporal relationships are utilized. Hippocampal time cells therefore serve as a cognitive map, representing temporal relationships in a manner that reflects cognitive demands.

## Introduction

The brain is thought to form cognitive maps of learned relationships within the world (Tolman 1948; O’Keefe & Nadel, 1978). Such maps have traditionally been studied in the hippocampus, where spatially tuned place cells provide an internal representation of an animal’s spatial environment (O’Keefe & Dostrovsky 1971). Although place cells fire when an animal occupies a specific region in space, multiple pieces of evidence demonstrate that place cells are not only tuned to position within an environment, but also reflect features of that environment. Most notably, place cells “remap”, forming distinct “maps” of unique environments (Muller & Kubie 1987; Colgin et al. 2008). Cell tuning is also dynamic within an environment; spatial maps change as animals gain exposure to and knowledge of a particular context. For instance, new place fields form as an animal spends more time in a context (Frank et al. 2004; Sheffield & Dombeck 2017), and place fields can sharpen their tuning to represent space more precisely (Roth et al. 2012). Place cells also strengthen representations of salient locations (Hollup et al. 2001; Dupret et al. 2010), reflecting their learned significance. These findings demonstrate that spatial tuning updates dynamically to reflect familiarity with or knowledge of a spatial context.

In addition to learning the spatial relationships that define a context, animals must learn how to use those relationships to guide behavior. This problem is apparent in tasks that require animals to change behavior systematically across trials. For instance, spatial alternation tasks require animals to alternate routes across trials. Remarkably, in such tasks, hippocampal “splitter cells” display differential activity in the same spatial location based upon which side of the maze the animal previously visited (Frank et al. 2000; Ferbinteanu & Shapiro 2003; Wood et al. 2000; Hasselmo & Eichenbaum 2005). In another task where lap number was behaviorally relevant, hippocampal activity tracked lap number (Sun et al. 2020). These findings demonstrate that hippocampal maps not only reflect knowledge of the spatial relationships within a context, but also reflect knowledge of broader task structures that dictate how spatial relationships should be used to guide behavior.

Both space and time are organizing features of experience, raising the question of whether the hippocampus similarly uses “cognitive maps” to represent temporal features of experiences. Just as hippocampal place cells represent a particular location in space, hippocampal time cells fire at a specific moment in a temporally structured experience (Pastalkova et al. 2008; MacDonald et al. 2011; MacDonald et al. 2013; Sabariego et al. 2019; Shimbo et al. 2021; Salz et al. 2016; Taxidis et al. 2020). Hippocampal time cells are thought to map the temporal structure of experiences (Eichenbaum 2014). However, while the temporal structure of experiences is complex, consisting of durations of and between multiple events, time cells are typically studied during one timed period of behavioral tasks (MacDonald et al. 2011; Sabariego et al. 2019; Shimbo et al. 2021; Salz et al. 2016). Therefore, it remains unclear whether or how the hippocampus represents temporal contexts consisting of distinct patterns of stimulus durations. Additionally, it is unknown whether hippocampal time cells display learning dynamics that reflect an increased knowledge of the temporal relationships that define a context, as well as how to use those relationships to guide behavior within a broader task structure.

To address this gap in knowledge, we asked how the hippocampus represents temporal context, and whether representations change with learning. We tested three predictions. First, if hippocampal time cells encode the temporal structure of experience, they should represent context from the first moments of exposure and should differentially encode distinct contexts. Second, we predicted that time cell tuning would change with experience, reflecting increased familiarity with contexts. Finally, we tested whether contexts are encoded in a manner that reflects behavioral or cognitive demands. Evidence of these features would suggest that hippocampal time cells serve as a cognitive map, representing knowledge of the temporal structure of experiences. The key to our study design was our behavioral approach: we used a curriculum-based approach that progressed from simple shaping phases to a complex timing task, and systematically observed how neural dynamics and behavior evolve as mice build up knowledge of temporal relationships. Consistent with our hypothesis, we find that CA1 time fields emerge and develop more precise fields with experience. By comparing activity across contexts, we find that distinct patterns of time cell activity tile each context. Additionally, the way contexts are represented relative to each other changes with experience: contexts are represented similarly when mice can generalize across contexts, and distinctly when mice must discriminate. Our data show that CA1 time cells not only encode temporal context, but also reflect knowledge of context in a behaviorally relevant way. Therefore, hippocampal time cells form a cognitive map, representing temporal relationships in a manner that reflects knowledge of how those relationships should be used.

## Results

### CA1 time cells are present at each training phase

To study how learning affects representations of temporal context, we trained mice on our temporal Delayed Non-Match to Sample (tDNMS) task (Bigus et al. 2024) (Figure 1A). In each trial, a head-fixed mouse is presented with two odor stimuli separated by a brief (3s) interstimulus interval (ISI) and trained to report if stimuli differ. The same odor is used each presentation so that stimulus identity is constant. Instead, the relevant stimulus feature is duration, with odors being either short (2s) or long (5s). The tDNMS task has 3 trial types: Long-Short (LS), Short-Long (SL), and Short-Short (SS), defined by the series of durations that form each trial. In a Go/No-Go fashion, mice are trained to lick to report a nonmatch of durations (LS & SL trials) and withhold in response to matched durations (SS trials). Due to the unique combinations of stimulus durations, each trial type has a unique temporal structure, or temporal context.

**Figure 1.**
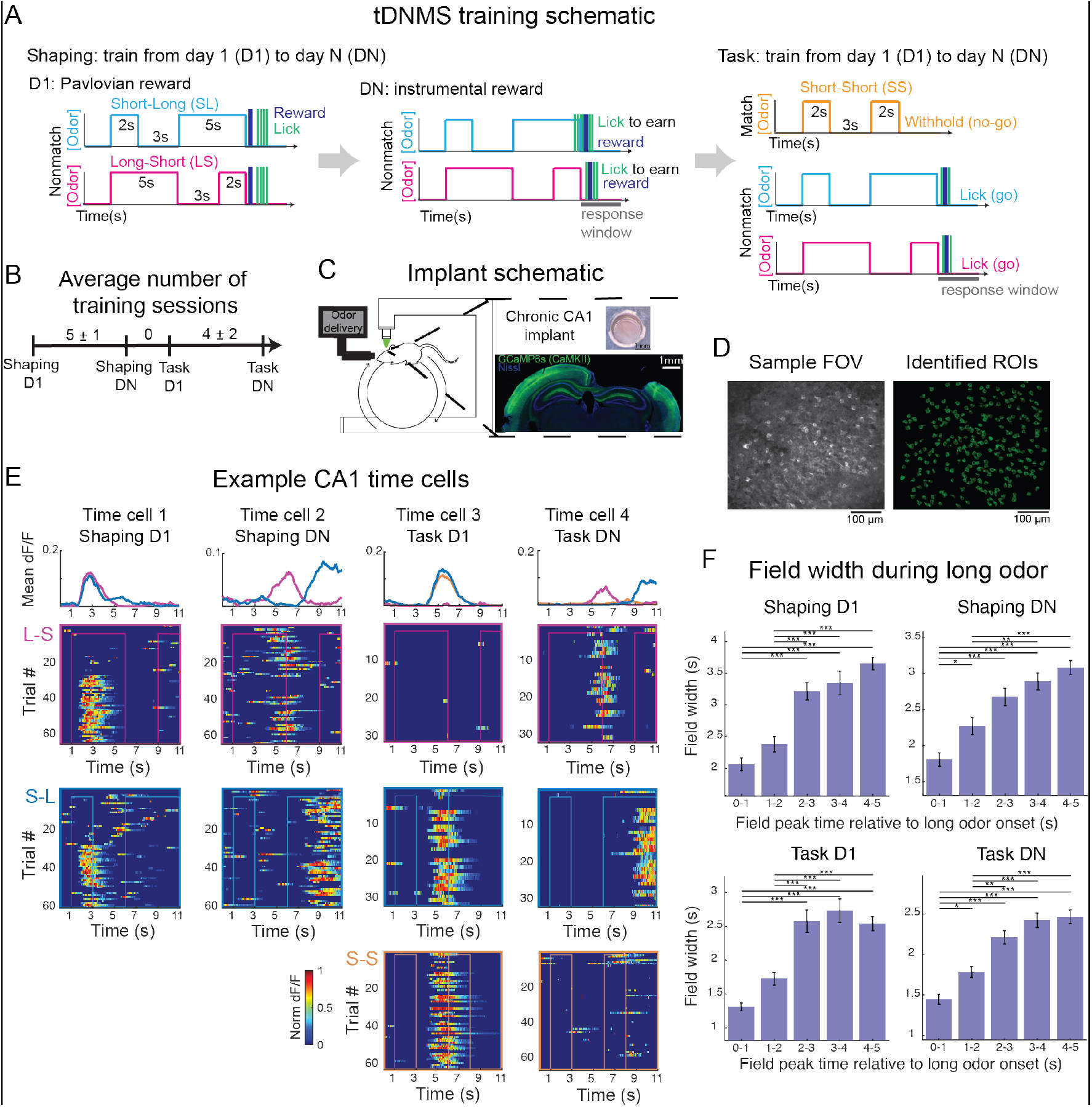
Time cells are present at each training phase. **A**. Training paradigm. Mice are trained from day 1 (D1) to day N (DN) on Shaping, then from day 1 (D1) to day N (DN) on the tDNMS Task. In shaping, mice progress from Pavlovian to instrumental training to learn the trial structure “odor-odor-respond”. In the task, mice are trained in a go/no-go manner to withhold licking if odor durations are matched. **B**. Average number of sessions between each training phase, shown as mean ± standard deviation. **C**. Implant schematic. Left-schematic of head-fixed mouse. Top right-example implant. Bottom right-histology demonstrating implant location. **D**. Sample field of view (FOV) and identified cells (regions of interest, ROI). **E**. Sample CA1 time cells. One cell is shown from each training phase (from left to right: Shaping D1, Shaping DN, Task D1, Task DN). Activity is shown for all trials, separated by trial type (from top to bottom: LS, SL, SS). Mean dF/F is plotted for each context (top). **F**. Field width during long odor stimuli. Data were combined across mice (Shaping D1 n = 4; Shaping DN n = 4; Task D1 n = 7; Task DN n = 8 mice) and include time fields with peaks during the long odor in LS and/or SL trials. Bars indicate mean field width ± SEM. Field width increases through long odor stimuli (p = ≤ 0.001 for each plot; ANOVA with Bonferroni post-hoc testing: *p ≤ 0.05, **p ≤ 0.01, ***p ≤ 0.001).

To aid the learning process, mice undergo several training phases (Figure 1A-B). Training begins with shaping, where the goal is to teach mice the trial structure: “odor-odor-response”. During shaping, only nonmatch (LS & SL) trials are presented. On the first day of shaping (Shaping D1), the task is Pavlovian, with reward automatically delivered at the end of each trial. As shaping progresses, training becomes instrumental, requiring mice to lick selectively at the second odor offset to earn a reward. By the end of shaping (day N), mice must withhold licking during the first odor and ISI and lick at second odor offset to earn reward, indicating awareness of the “odor-odor-response” trial structure. During shaping, odor duration is not relevant, as mice only need to respond at the second odor offset. After completing shaping (Shaping DN), mice begin the tDNMS task, which introduces matched trials where licking at the trial’s end is not rewarded. The inclusion of matched trials changes the task into a timing task, requiring mice to use stimulus durations to determine whether to lick or withhold on each trial. Mice are trained on the tDNMS task from day 1 (task D1) until post-learning day N (task DN). To study hippocampal activity across learning, we used established methods for cellular-resolution two photon imaging in CA1 (Dombeck et al. 2010) (Figure 1C-D). We trained mice to perform the tDNMS task and imaged at the 4 described phases of training: Shaping D1 (n = 4 mice), Shaping DN (n = 4 mice), Task D1 (n = 7 mice), Task DN (n = 8 mice).

If CA1 time cells represent the temporal structure of experiences, time cells should meet several predictions. First, time cells should tile any experience and thus be present at each training phase. Second, if time cells reflect an internal timing process, a key prediction is that their dynamics should exhibit scalar properties (Gibbon 1977). To test these predictions, we first identified time cells (Figure 1E). Consistent with our first prediction, a large proportion of cells displayed time locked activity in each training phase: approximately 54%, 59%, 39%, and 53% of cells were classified as time cells on Shaping D1, Shaping DN, Task D1, and Task DN respectively (Shaping D1: 540/1009 cells from 4 mice; Shaping DN: 555/940 cells from 4 mice; Task D1: 714 of 1809 cells from 7 mice; Task DN: 1001 of 1898 cells from 8 mice). Next, we examined whether the scalar property was maintained (Figure 1F). We divided each long odor period into 1-second intervals, or bins, and identified time fields with peaks in each bin (Figure S1). Then, we calculated the average width of all time fields with peaks in each bin to assess how field width changes over time. In each training session, we observed that field width progressively increased across the long odor duration (p ≤ 0.001 for each session), suggesting that time cells may be tracking time in a manner consistent with scalar timing properties.

### Time fields become more precise with experience

If time cells map experiences, they should be present from the first trial, not just on the first training session. To test this prediction, we examined the activity of cells on Shaping D1 (Figure 2A). As expected, some time cells (21%) were active from the very first trial. Interestingly, others formed fields later in the training session. We noticed that time cells also displayed differences in other field properties, including number of fields, field width, and field stability. Certain properties, like having fewer or narrower fields, provides more precise information. Therefore, we wondered whether cell tuning properties change across training sessions, perhaps reflecting increased familiarity with temporal contexts.

**Figure 2.**
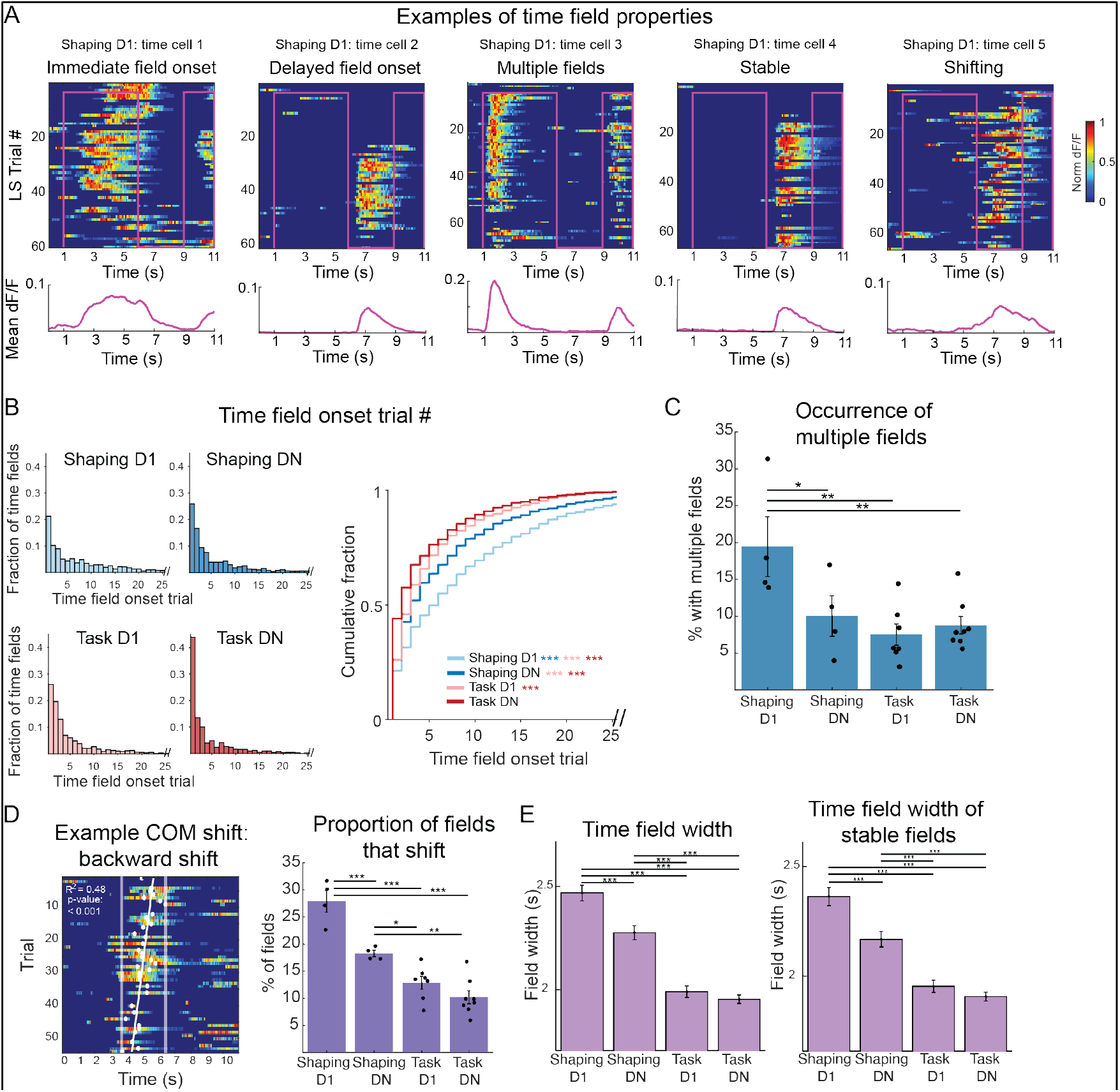
Time field proper8es change with training. **A**. Sample Gme cells on Shaping D1. The acGvity of each cell is shown for each LS trial (top). Mean dF/F is ploPed below. Time fields display different properGes, noted above plots. **B**. Time field onset trial. LeL-histograms showing the fracGon of fields that emerged on each trial number of a context. Analysis includes all idenGfied fields on each session (n = 1037, 988, 1327, 2104 fields for Shaping D1, Shaping DN, Task D1, Task DN respecGvely). Right-cumulaGve fracGon plot comparing distribuGons across each training phase. Fields emerge earlier across training phases (p = 1.41e-68; Kruskal-Wallis test with Dunn-Bonferroni post-hoc tesGng: *p ≤ 0.05, **p ≤ 0.01, ***p ≤ 0.001 in all plots). **C**. Occurrence of mulGple fields. For each cell x context pair with at least 1 Gme field, it was determined whether there was 1 field or mulGple. This calculaGon was done for all fields from a given mouse. Data points represent proporGons from individual mice, and bars show mean across mice ± SEM. Occurrence of mulGple fields decreases over sessions (p = 0.0024; linear mixed effects model). **D**. LeL-example of center of mass (COM) shiL. Rightpropor Gon of fields that shiLed on each session. Dots show proporGons for individual mice, and bars indicate mean across mice ± SEM. Across training, the proporGon of shiLing fields decreases (p= 3.20e-08; linear mixed effects model). **E**. LeL-average Gme field width per session, shown as mean ± SEM. Field width decreases with training (ANOVA p = 9.89e-46 with Bonferroni post-hoc tesGng *p ≤ 0.05, **p ≤ 0.01, ***p ≤ 0.001 for all figures). Right-average Gme field width only including stable (non-shiLing) fields ± SEM. Width of stable fields decreases with training (ANOVA p = 1.07e-31 with Bonferroni post-hoc tesGng).

We began by quantifying field onset (Dong et al. 2021) (Figure 2B), hypothesizing that with experience, fields should emerge earlier, indicating increased familiarity of trial structure. Indeed, across training sessions, time fields emerge in earlier trials (p ≤ 0.001). Next, we asked if field number (Figure 2C) changes with experience, perhaps decreasing to become more precise. For each mouse, we identified all cell x context pairs with at least one time field, then calculated the proportion with multiple fields. As predicted, the occurrence of multiple fields decreases across sessions (p = 0.0024).

Spatial literature has described how place fields can display shifts in their center of mass (COM) (Mehta et al. 1997), which is thought to be driven by behavioral timescale synaptic plasticity (BTSP) and contribute to the formation of spatial maps (Bittner et al. 2017; Grienberger & Magee 2022). Therefore, we examined whether time fields likewise shift (Figure 2D). For this analysis, we found each field’s center of mass on each trial (of a specific context) then used linear regression to determine whether the field significantly shifted over the course of the session (Dong et al. 2021). Not only did fields shift, but relatively fewer fields shifted later in training, indicating increased field stability (from 27.9% of fields on Shaping D1 to 10.2% on Task DN; p ≤ 0.001).

Finally, though we cannot quantify absolute field width due to the time course of our calcium indicator, we noticed that fields displayed differences in relative widths. We asked if field width decreases with training, making time fields more precise (Figure 2E). Indeed, field width significantly decreases (p ≤ 0.001). One factor that influences field width is stability; fields with a more stable center of mass may be narrower than fields with a shifting center of mass. One explanation for the observed decrease in field width is that fields are simply more stable (Figure 2D). Therefore, we repeated field width analysis, this time only on stable fields. Field width still decreased, showing this is an experience-dependent effect (p ≤ 0.001).

### Is time cell activity better explained by velocity

Though time cells display changes in activity in time, like field shifting, an alternative possibility is that these changes can be explained by other factors, like changes in mouse velocity. Given the strong spatial tuning in the hippocampus, we therefore asked whether changes in time cell properties, specifically field shifting, were due to changes in velocity (Figure S2). While some field shifts were indeed correlated with velocity (Figure S2F), we observed the same pattern in field shifts across sessions even excluding such cells (Figure S2 G-I), suggesting changes are not driven by velocity.

Further, before interpreting behavioral data, we wanted to rule out the possibility mice used a distance-tracking strategy to solve the task. Although the task is performed in darkness without visual cues, mice could use distance travelled to estimate when they should lick. If this were the case, mice should lick earlier on trials with higher velocity, as they reached a putative distance goal earlier. This is not the case (Figure S2B-C), indicating that mice do not use a distance tracking strategy to solve the task.

Finally, to confirm that time cells are tuned to time, we calculated the coefficient of variation of time cell activity (CV) when measured as a function of time compared to distance (Figure S2D). The CV is smaller for time than distance, giving confidence that identified cells track time not distance.

### Distinct patterns of time cell activity represent each temporal context

If CA1 time cells represent the temporal structure of experiences, cells should meet two additional predictions. First, time cells should span the full duration of each trial. Second, distinct patterns of time cell activity should represent distinct temporal experiences, or contexts. By plotting the trial averaged activity of time cells, we first confirmed that time cells tile the entire trial period (Figure 3A). By sorting cells based on their peak time of activity in SL trials, then applying that same sorting to other contexts, we could also see that time cell sequences appear to differ across contexts. Though additional quantification, we confirmed that sequences differ soon after stimuli diverge (Figure S3D-F), demonstrating that representations of context are distinct.

**Figure 3.**
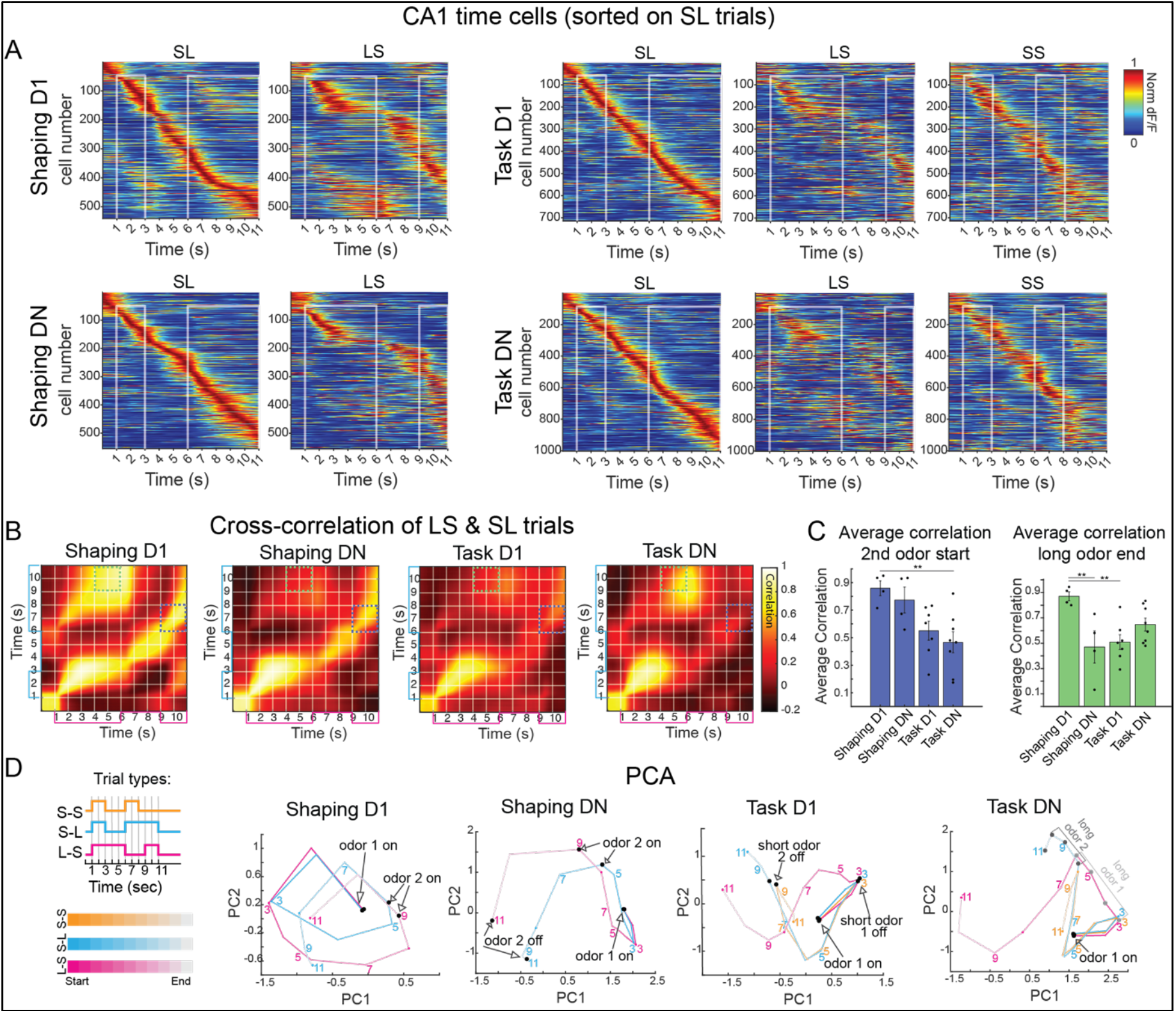
Relative representations of context change across training phases. **A**. Population of CA1 time cells sorted by response times during SL trials and shown for other contexts. Plots are shown for Shaping D1 (top left), Shaping DN (bottom left), Task D1 (top right), and Task DN (bottom right). **B**. Population vector cross correlation matrices, shown for LS & SL trial types across each training phase (from left to right-Shaping D1, Shaping DN, Task D1, Task DN). **C**. Population vector correlations were averaged during key trial periods, specifically the first 2s of each 2^nd^ odor (left) and the last 2s of each long odor (right). Dots show values for individual mice, and bars represent mean ± SEM across mice. Across sessions, there is a significant difference in correlations during both the 2^nd^ odor start (p = 0.004; linear mixed effects model with post hoc pairwise comparisons *p ≤ 0.05, **p ≤ 0.01) and long odor end (p = 0.0032; linear mixed effects model). **D**. For each session, PCA was performed using time cell data. Trajectories of each trial type are plotted on the first 2 principal components. Numbers indicate time (s) in the trial.

### Similarity or difference between contexts align with learning demands

Just as single cell tuning changes with experience, we hypothesized that population-level representations of contexts may also change with experience. We first asked whether representations of context gradually become more or less similar with each phase of training. However, by comparing the correlation of time cell activity across contexts for each session, it is clear this is not the case (Figure S3A), suggesting representations of context may vary in a more nuanced manner.

Although SL and LS contexts are present in each training phase, with training, mice may develop a better understanding of how to use these contexts to guide their behavior. Additionally, the demands of each training phase differ: in the shaping phase, mice do not need to discriminate between temporal contexts, whereas in the tDNMS task, context discrimination becomes essential. We therefore questioned whether contexts become increasingly dissimilar throughout training, particularly at key task moments when mice can differential context, reflecting increased awareness of the differences in contexts and knowledge of how to use contexts.

To examine similarity between contexts, we created population vectors from trial averaged time cell activity at each moment in SL and LS contexts. We then plotted the correlation of population vectors in a cross-correlation matrix (Figure 3B and S5). We noticed striking differences in cross-correlation matrixes across training sessions. First, on Shaping D1, activity is highly correlated during each long odor (Figure 3C). This correlation decreases on Shaping DN, but activity remains highly correlated at the same relative phases in each trial, for instance, the start of the second odor (Figure 3C) Compared to Shaping, in the Task, activity is less correlated at relative task phases like 2nd odor start (Figure 3C). Instead, there is a strong on-diagonal correlation in the beginning of each trial, when stimuli are identical, which decreases soon after stimuli diverge. Task learning is further characterized by an additional off-diagonal correlation during the long odor period (Figure 3B). These differences across sessions indicate that the ways contexts are encoded relative to each other changes throughout the training process. Each phase of training has distinct cognitive demands. Therefore, we wondered whether the way contexts are encoded aligns with behavioral demands, consistent with our overarching hypothesis that time cells serve as a cognitive map, reflecting knowledge of each temporal context and its use in a larger task structure.

We subsequently reflected on the learning requirements in each training phase. On Shaping D1, reward is Pavlovian, meaning mice do not need to attend to trial structure to earn reward (Figure 1A). Accordingly, mice may initially recognize individual odors and pauses but might not yet understand the two-odor trial structure. If this is the case, we would expect similar neural activity during both the first and second odor periods within each trial. Indeed, upon reexamining trial-averaged time cell sequences (Figure 3A), it is clear that the same cells are often active during odor presentations in both SL and LS trials.

By Shaping DN, training becomes instrumental, and mice must lick selectively near second odor offset to earn reward (Figure 1A). This behavior requires an awareness of the “odor-odor-respond” trial structure. At this training phase, mice can generalize across contexts by consistently following an ‘odor-odor-response’ strategy. This generalization is reflected in the neural activity (Figure 3A): the same cells tend to activate at corresponding task phases—such as the start and end of the ISI and the onset of the second odor—in both SL and LS trials (Figure S4). Accordingly, there are high correlations in time cell activity between SL and LS trials at these analogous phases, even though they occur at different absolute times (Figure 3B).

In Task D1, matched trials are introduced (Figure 1A), and mice are no longer rewarded for licking at each trial end. Remarkably, when it is no longer useful for mice to generalize across trials, time cells tend to lose fields in one context (Figure S4C), no longer representing each context (Figure 3A) Instead, population activity is only highly correlated during the initial trial period when contexts are identical (Figure 3B).

From Task D1 to Task DN, mice gain experience with SS trials. Because shaping only includes SL and LS trials, an ideal observer could discriminate trial type only using the first duration. Given the extensive pretraining process, we suspect that mice were able to discriminate SL and LS contexts by the first odor duration, even before this was a behavioral requirement. The addition of SS trials in the Task requires mice to use the second odor duration to discriminate SS and SL trials. Perhaps as mice grow accustomed to discriminating to second odor duration, they begin to use similar neural dynamics to differentiate both the first and second odor as being long, explaining the Task DN correlation in the long odors of SL and LS contexts (Figure 3B).

Together, our data demonstrate that contexts are not encoded in the same manner across all training phases. Rather, the way contexts are encoded relative to each other changes with behavioral demands, with contexts encoded similarly when mice can generalize and distinctly when mice must discriminate context. We next asked if we could find additional support that the way contexts are encoded changes with cognitive demands. Therefore, we performed PCA on trial-averaged activity of time cells and plotted resulting trajectories for each trial type (Figure 3D). As anticipated, PCA results complement our prior analysis. On Shaping D1, SL and LS trajectories display similar patterns during the first odor, and at second odor onset, trajectories loop back to the location of first odor onset, reflecting similarity in neural dynamics during each odor presentation. By Shaping DN, trajectories no longer loop and instead display a less tangled trajectory spanning the trial. The shape of LS and SL trajectories is remarkably similar, with the same relative moments in each trial epoch lying in similar places in state space, confirming that neural activity is similar during analogous trial phases. While SL and LS trajectories follow a similar path in Shaping DN, this is no longer the case on Task D1, as trajectories split in different directions soon after stimuli diverge, confirming contexts are represented differentially. Finally, we looked for evidence that in Task DN, mice apply similar dynamics to distinguish both the first and second odor duration. On Task DN, but not previous sessions, activity during the timepoints that allow animals to discriminate each odor as being long fall on the same axis in our PCA plot, confirming commonality in neural dynamics during these classification periods.

Despite SL and LS contexts remaining identical across training, the way these contexts are represented relative to each other changes. Strikingly, the way contexts are encoded offers insights into how mice may think about context in each training phase. Specifically, time cell dynamics suggest that from Shaping D1 to Shaping DN, mice learn the “odor-odor-respond” trial structure. Mice seem to generalize across contexts in shaping, encoding relative moments of each trial similarly. In contrast, when mice need to discriminate context in the tDNMS task, dynamics become increasingly distinct across contexts. Like our single cell analysis, these results show that time cell activity changes in an experience dependent manner. More specifically, time cell activity reflects knowledge of how mice should use each context, consistent with time cells providing a cognitive representation that reflects salient features of temporal context.

### Behavior changes across training phases

Temporal contexts are represented differently across training, suggesting animals may think about or use contexts distinctly at each training phase. We therefore asked whether there was behavioral evidence that mice use contexts differently across training. To examine behavior, we began by plotting licking behavior on each trial (Figure 4A). If mice encode odors similarly on Shaping D1, behavior may be similar during each odor presentation, regardless of whether it is the first or second odor in a trial. Indeed, in Shaping D1, even though only the second odor is rewarded, mice also lick at first odor offset in 37.5% of trials, as if treating odors similarly (Figure 4B). With further training, mice learn to withhold licking after the first odor (p = 0.0027) and increase predictive licking in the second odor (p = 0.0053). This combination of learning to withhold licking in the first odor and ISI, and lick in the second odor and response window of nonmatch trials, leads to mice displaying increasingly distinct licking behavior across odor 1 and 2 over sessions (p = 0.0059). Additionally, while mice reliably lick at second odor offset in the majority of trials in Shaping D1 (86.5%) and DN (93.6%), mice quickly learn to apply a new strategy on Task D1 and begin withholding licking, only licking in the response window of 60.3% of trials. This change in licking in the response window across sessions (p = 8.99e-12) is a result of the introduction of match trials and strong performance in this trial type (Figure 4E). By withholding licking in SS and maintaining high performance in other trial types (Figure 4D), mice no longer treat each context similarly. By Task DN, mice have gained experience discriminating odor durations and show improved task performance (Figure 4E, p = 0.0019), driven by an improvement in SS trials (p = 0.0022). Therefore, transitions in training phases are accompanied by behavioral changes.

**Figure 4.**
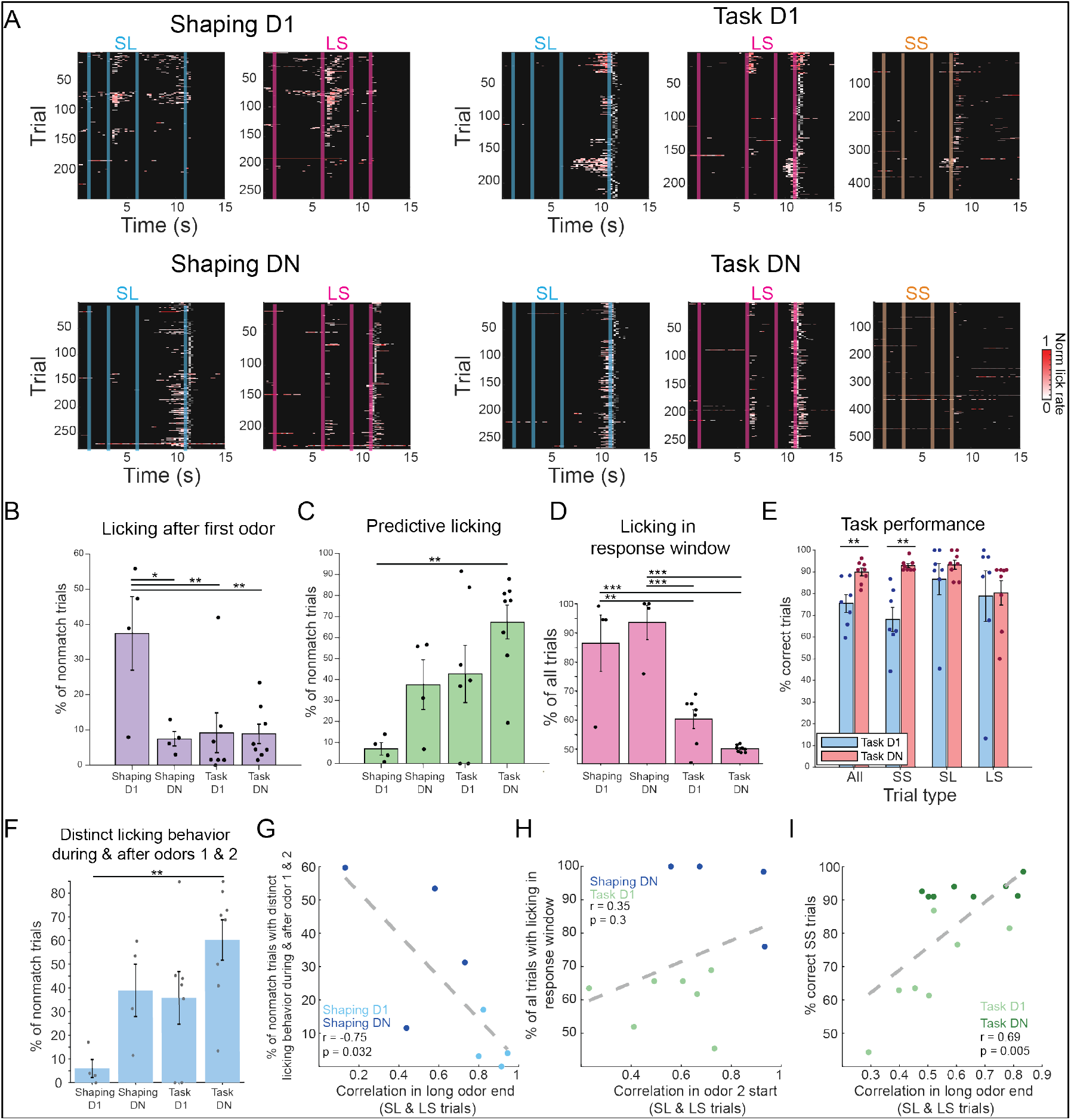
Mouse behavior changes with training. **A**. Licking behavior, shown for each trial for each mouse on imaging sessions (Shaping D1 n = 4; Shaping DN n = 4; Task D1 n = 7; Task DN n = 8 mice). Trials are separated by trial type. Licking following reward delivery is not shown. **B**. Percent of nonmatch trials with licking in the 3s interstimulus interval (ISI) after the first odor. In all bar plots, dots indicate values per mouse, with bars showing mean across mice ± SEM. With training, mice learn to withhold licking after odor 1 (p = 0.0027; linear mixed effects model with post hoc pairwise comparisons *p ≤ 0.05, **p ≤ 0.01, *** p ≤ 0.001). **C**. Percent of nonmatched trials with predictive licking, defined as licking in the last 0.5s of the second odor. Predictive licking significantly increases with training (p = 0.0053; linear mixed effects model). **D**. Percent of all trials with licking in the 3s response window following 2nd odor offset (p = 8.99e-12; linear mixed effects model). **E**. Task performance on Task D1 (blue) and Task DN (red). Performance is shown across all trials and by trial type and is significantly different between D1 and DN for all (p = 0.0019) and for SS trials (p = 0.0022, paired t-test). **F**. Percent of nonmatch trials with distinct licking behavior during and after odors 1 and 2. Distinct licking behavior is defined as trials with no licking during odor 1 and in the 3s ISI after, and with licking during odor 2 and in the 3s response window after (p = 0.0059; linear mixed effects model). **G.** As the correlation in long odor end (from the cross correlation matrixes in Figure 3B-C) decreases, mice display more distinct licking behavior during and after odors 1 and 2, as defined in F (p = 0.032; Pearson’s correlation). Dots show values per mouse, with light blue for Shaping D1 and dark blue for Shaping DN. **H.** There is no significant relationship between the correlation in odor 2 start (Figure 3B-C) and percent of all trials with licking in the response window (D) from Shaping DN (dark blue) to Task D1 (light green) (p = 0.3; Pearson’s correlation). **I.** Correlation in long odor end (Figure 3B-C) on Task D1 (light green) and N (dark green) is correlated with task performance on SS trials (E) (p = 0.005; Pearson’s

Finally, to test if changes in neural activity may support changes in behavior, we examined neural activity and behavior of individual mice. First, we predicted that if having similar representations of long odors in Shaping D1 compared to DN caused mice to treat odors more similarly, sessions with higher correlations (Figure 4C) should have more similar licking behavior. Indeed, this is the case (Figure 4G; p = 0.032). Next, we asked whether a lower correlation in the second odor (Figure 4C) might help mice treat contexts more distinctly in Task D1 compared to Shaping DN. While this was not the case (Figure 4H; p = 0.3), the correlation in long odor end is correlated with performance on SS trials (p = 0.005; Figure 4I), suggesting the increased similarity of long odors helps mice solve the task.

## Discussion

The hippocampus is known to form cognitive maps—internal representations of relationships within the world (Tolman, 1948; O’Keefe & Nadel, 1978). Although hippocampal research has traditionally focused on spatial maps, we designed this study to explore whether and how the hippocampus builds knowledge of temporal context through a structured learning curriculum. Using a curriculum-based approach that progressed from simple shaping phases to a fully complex task, we systematically observed how neural dynamics and behavior evolve as mice build up knowledge of temporal relationships. We tested the prediction that hippocampal time cells would display learning dynamics reflecting this stepwise acquisition of temporal context. While time cells were present from the first moments of training, fields emerged and became more informative with experience. Importantly, we also found that time cell activity patterns reflected a cognitive component: contexts were represented similarly when mice could generalize and distinctly when they had to discriminate. This structured approach demonstrates that time cells not only represent temporal context but also adapt with cognitive demands, supporting a more nuanced view of cognitive mapping in the hippocampus.

Two key bodies of work demonstrated that spatially tuned cells in the hippocampus form a cognitive map: 1) place cell tuning changes with familiarity in a particular context and 2) neural codes reflect how spatial information should be used to meet cognitive or behavioral demands. We not only show that CA1 time cells represent temporal context, but likewise demonstrate that temporal tuning 1) changes with experience in contexts and 2) undergoes population-level changes to encode contexts in a behaviorally relevant manner. These similarities in spatial and temporal coding contribute to a growing body of work demonstrating that medial temporal lobe regions form internal representations of both spatial and nonspatial relationships (Tavares et al. 2015; Theves et al. 2020; Neupane et al. 2024). In addition to the main parallels described above, many of our specific findings match observations from spatial literature. Like place cells, CA1 time cells can exhibit multiple or single fields (Fenton et al. 2008), stable or unstable fields (Mehta et al. 1997), differences in field width (Dong et al. 2021), and immediate or delayed fields (Frank et al. 2004; Sheffield & Dombeck 2017). Additionally, both place and time cells display differential activity across contexts (Muller & Kubie 1987; Colgin et al. 2008), and further encode information in a way that can aid discrimination (Frank et al. 2000; Ferbinteanu & Shapiro 2003; Wood et al. 2000; Hasselmo & Eichenbaum 2005) or generalization (Sun et al. 2020) across trials or contexts, according to task demands. Given these similarities, a key future direction will involve testing whether similar cells or mechanisms encode spatial and temporal information in the hippocampus.

Finally, while the strength of this study lies in its description of learning dynamics in CA1 time cells, we cannot be certain whether learning occurs locally or is inherited from upstream activity. Consistent with the latter, we previously identified learning dynamics in medial entorhinal cortex (MEC) time cells (Bigus et al. 2024). Though prior work suggests that CA1 time cells are not dependent on MEC activity (Sabariego et al. 2019), it remains unclear if this finding also holds true during explicit timing behavior. Our findings that both MEC and CA1 display learning dynamics in time cells pave the way for future studies to examine relative and potential differential contributions of each region to timing behavior. Together, our identification of learning dynamics in time cells strengthens a greater body of work, demonstrating that medial temporal lobe regions represent relevant features of experience, including time, in cognitive maps.

## Contributions

E.R.B, J.C.B and J.G.H. designed experiments and wrote the manuscript. E.R.B, J.C.B and D.Z. collected data. E.R.B, J.C.B, D.A. and J.G.H. analyzed and interpreted data. E.R.B. and J.G.H. built equipment used to collect data.

## Acknowledgments

We thank Matt Wachowiak for valuable comments on earlier versions of this manuscript and for generous support in designing and validating the olfactometers used in this study. This work was supported by the NIH/NIMH 1 DP2 MH129958-01, NSF CAREER Award: IOS-2145814, and the University of Utah.

---

**Figure S1.**
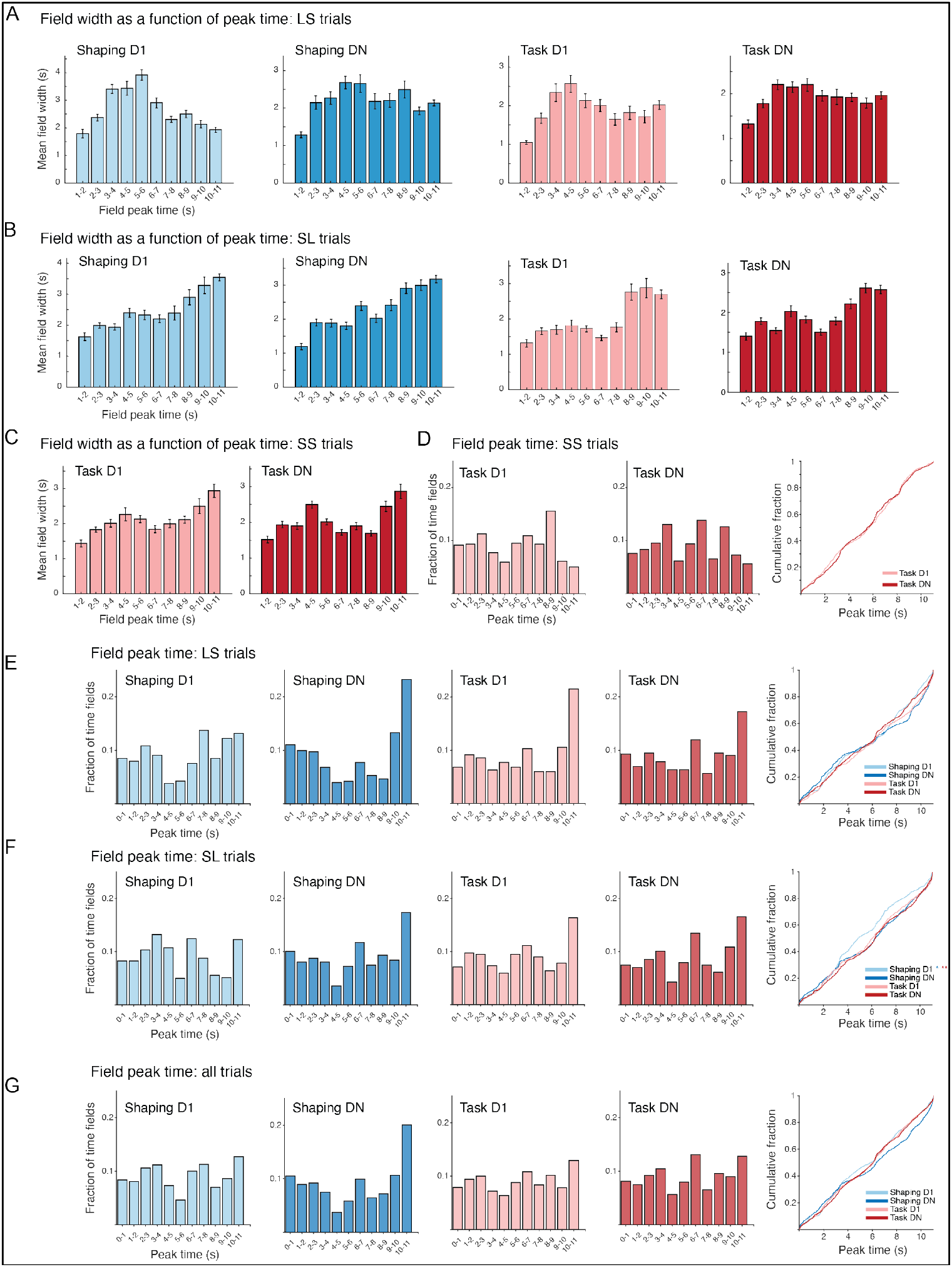
Additional analysis of time field peak and width. **A**. Time field width as a function of peak time for LS trials. Bars show average width of the time fields with a peak within each 1s bin. Bins span first odor onset (1s) to second odor offset. Histograms are shown for each session, and bars show mean ± SEM. **B**. Same analysis for SL trials. **C**. Same analysis for SS trials. **D**. Distribution of field peaks on SS trials. Left-histograms showing the fraction of time fields with a peak within each 1s bin of the trial, with first odor onset at 1s. Histograms are shown for each session. Far right-cumulative fraction plot comparing distributions across training phases (Kruskal-Wallis test with Dunn-Bonferroni post-hoc testing: *p ≤ 0.05, **p ≤ 0.01, ***p ≤ 0.001 in all plots). **E**. Same analysis for LS trials. **F**. Same analysis for SL trials (p = 0.0032). **G**. Same analysis for all trials.

**Figure S2.**
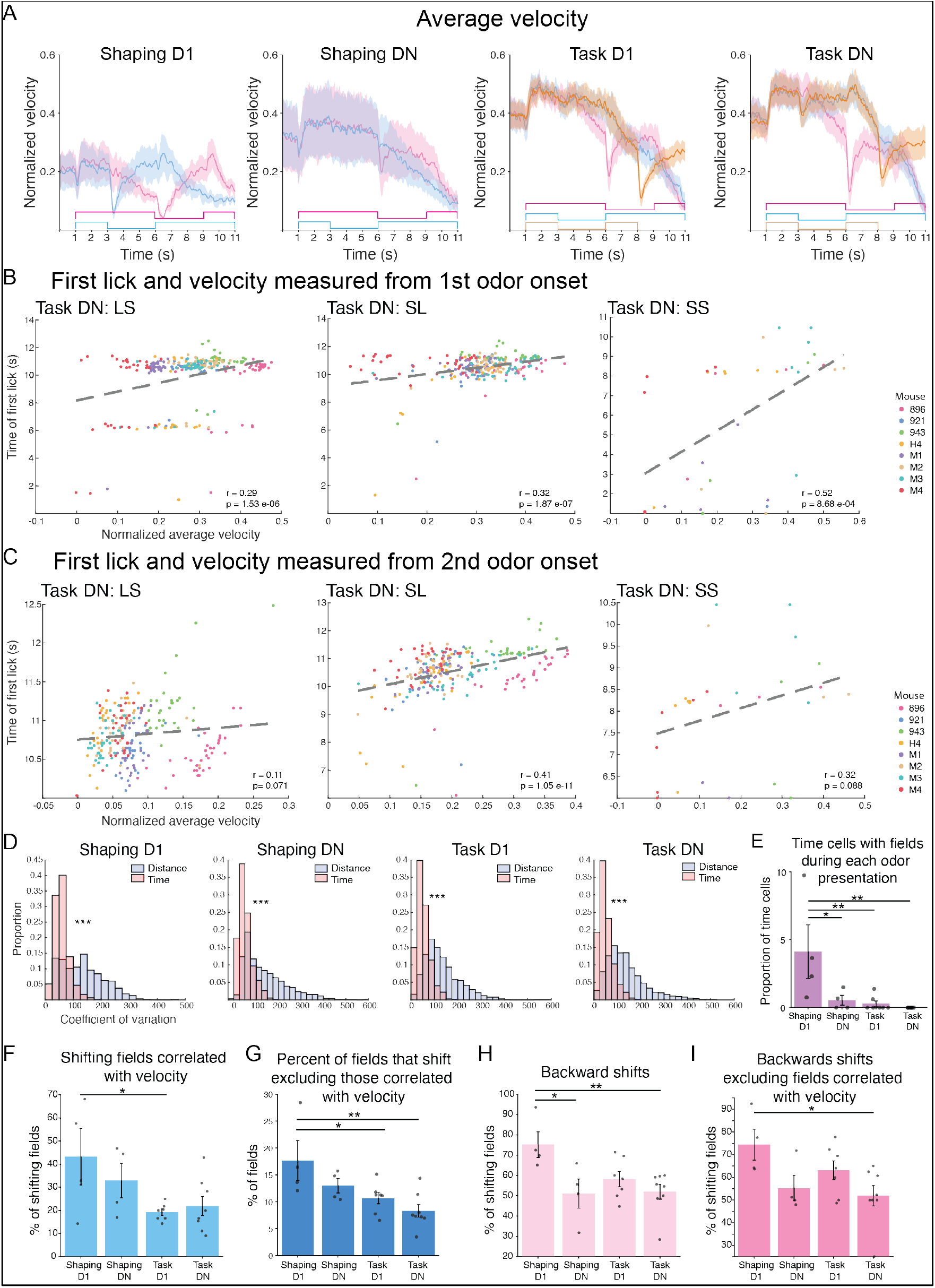
Time cell activity does simply reflect velocity or odor. **A**. Average velocity for each trial type and each session. Velocity was normalized by mouse to the maximum within the session. Lines depict average normalized velocity calculated across mice, and shading shows SEM (n = 4 Shaping D1, n = 4 Shaping DN, n = 7 Task D1, n = 8 Task DN). **B**. For each trial, the time of first lick and average velocity from first odor onset was measured. If mice track distance to solve the task, mice should lick earlier on trials with more running (higher velocity). However, across each trial type, the opposite relationship exists, where velocity is positive correlated with time of first lick (p < 0.001 for LS, SL, and SS trials). Each point represents a trial on Task DN, with dots colored to reflect mouse identity. **C**. Same as B, but with first lick and velocity measured from 2^nd^ odor onset (p = 0.071, 1.05e-11, 0.088 for LS, SL, SS trials respectively; Pearson’s correlation). **D**. Coefficient of variation measured as function of elapsed time or distance (from 1s before first odor onset). Distribution of values is shown for each session. In all sessions, the coefficient of variation is smaller when measured as a function of elapsed time (p < 0.001; paired t-test). **E**. Percent of time cells with a time field during each odor presentation in each context (p = 0.0013; linear mixed effects model with post hoc pairwise comparisons *p < 0.05, **p < 0.01). In all bar plots, dots show values for individual mice, with bars showing mean across mice ± SEM.**F**. Percent of shifting fields where field shift is correlated with velocity on a trial-by-trial basis (p = 0.022; linear mixed effects model) **G**. Percent of fields that shift, excluding all fields where shifts are correlated with velocity. Fewer fields shift with additional training (p = 0.0042; linear mixed effects model). **H**. Direction of field shift. The percent of shifts that are backwards (example in Figure 2D) decreases with training (p = 0.008; linear mixed effects model). **I**. Direction of field shift, excluding all fields where shift is correlated with velocity. The percent of backwards shifts decreases with training (p = 0.019; linear mixed effects model).

**Figure S3.**
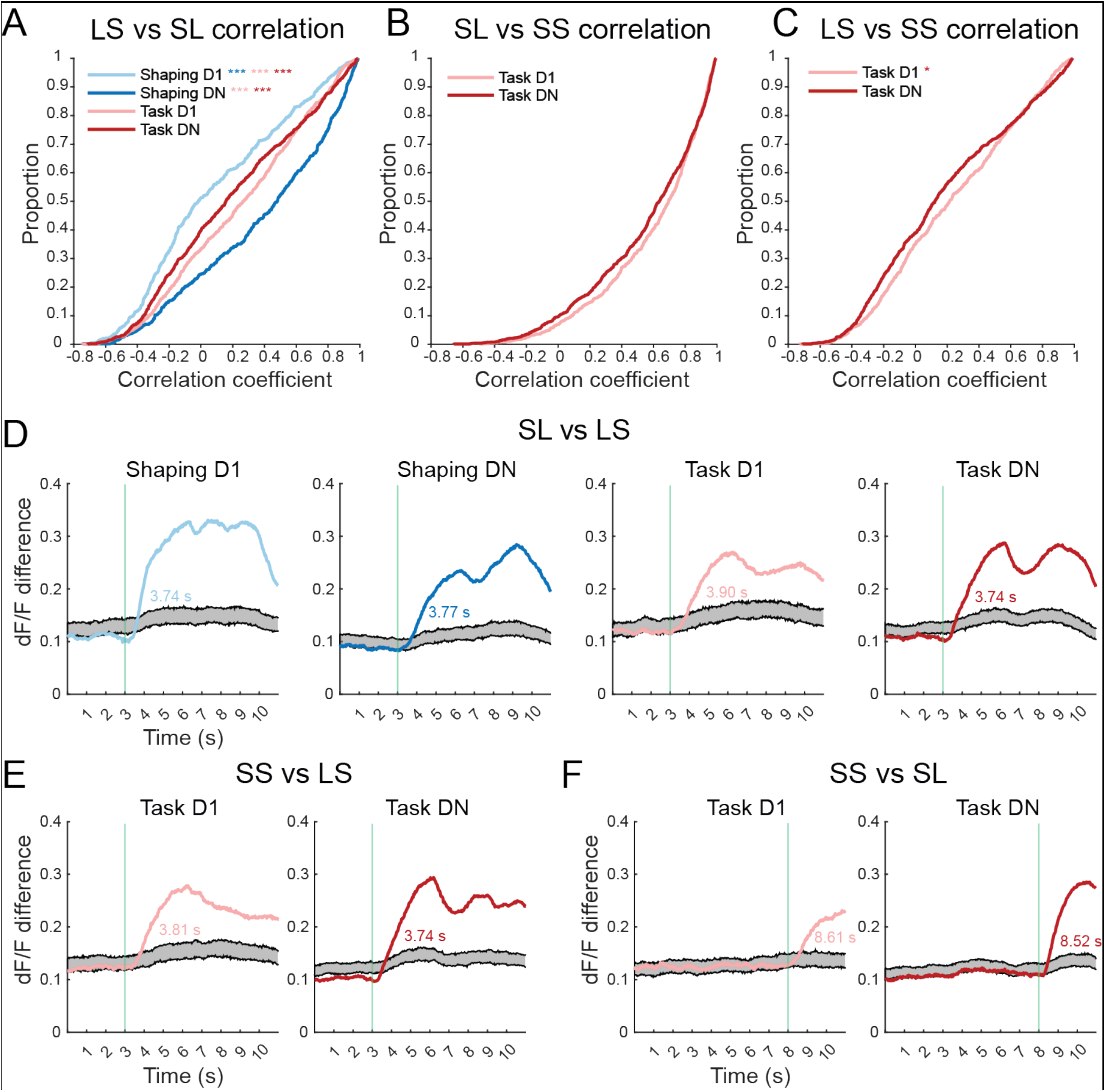
Representations of context are distinct. **A**. Average correlation of time cell rate maps (Pearson’s correlation) across LS and SL contexts, calculated for each session. Differences are significant across sessions (p = 7.53e-28; Kruskal-Wallis test with Dunn-Bonferroni post-hoc testing: *p ≤ 0.05, **p ≤ 0.01, ***p ≤ 0.001 in all plots). **B**. Same as A but for SL and SS contexts (p = 0.054; Wilcoxon rank sum). **C**. Same as A but for LS and SS contexts (p = 0.012; Wilcoxon rank sum). **D**. Population vector difference in dF/F for time cells between SL and LS trials at each moment in time, shown for each session. Colored lines represent true data, and gray lines represent shufled data distributions. Green bars demonstrate when stimuli diverge, and text indicates the time real data diverges from shuffled. **E**. Same as D but for SS and LS trials. **F**. Same as A but for SS and SL trials.

**Figure S4.**
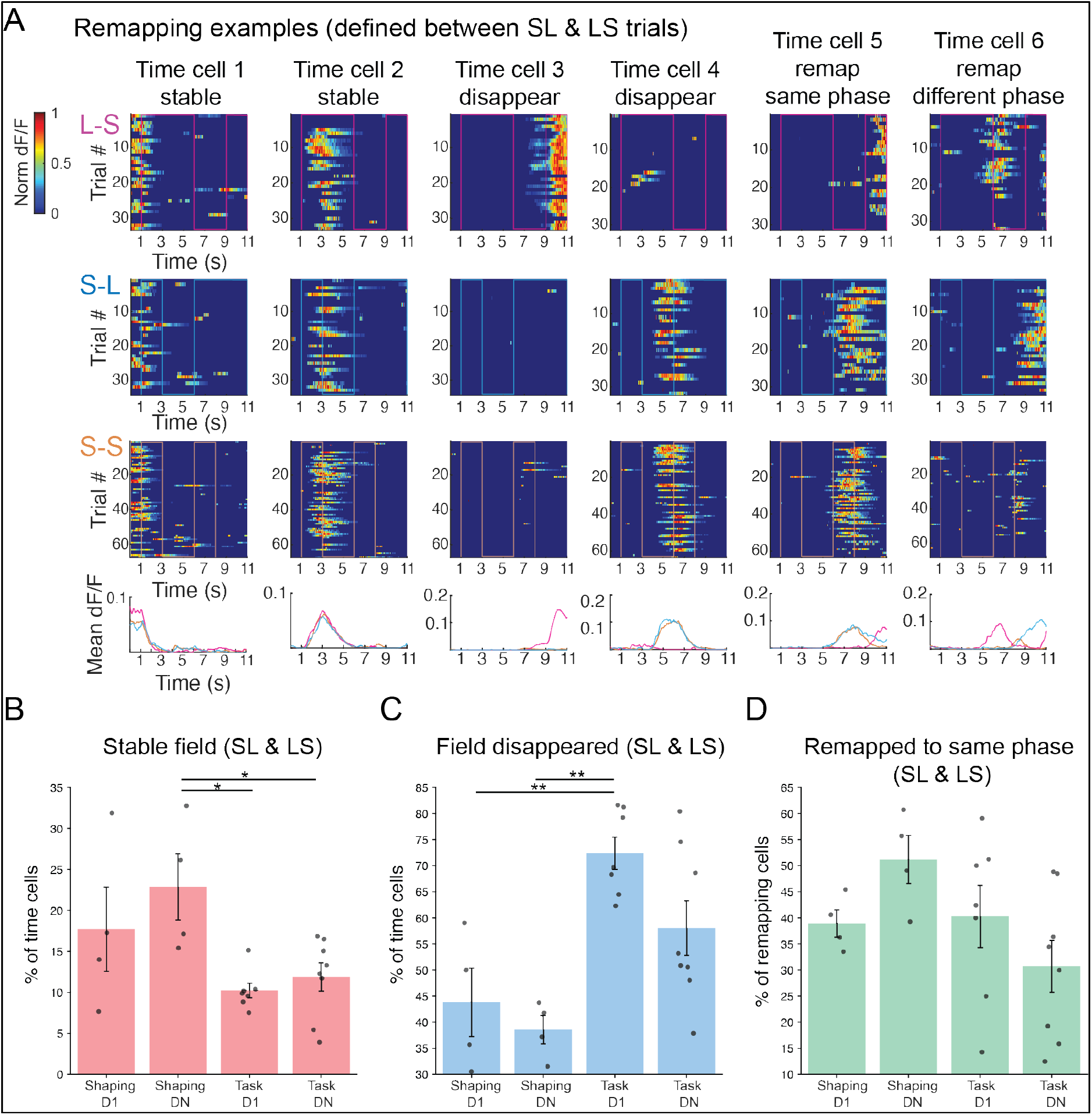
Time field remapping. **A**. Additional examples of time cells. Activity is shown for each trial, separated by trial type, with mean dF/F shown below. Time cell remapping was examined between LS (top) and SL (middle) contexts in this and subsequent plots. Some time fields are stable (cells 1 & 2), others disappear (cells 3 & 4) or remap (cells 5&6) across contexts. Fields can remap to the same task phase (for instance, odor 2-cell 5), or different task phases (cell 6). **B**. Percent of time cells with a stable field decreases over training (p = 0.0071; linear mixed effects model with post hoc testing *p ≤ 0.05, **p ≤ 0.01). In all plots, data points represent percent of cells per mouse, with bars showing mean ± SEM across mice. **C**. Percent of time cells in which a field disappears between contexts increases with training (p = 7.06e-04; linear mixed effects model). **D**. The percent of remapping cells that remap to the same task phase does not significantly change across training (p = 0.07; linear mixed effects model).

**Figure S5.**
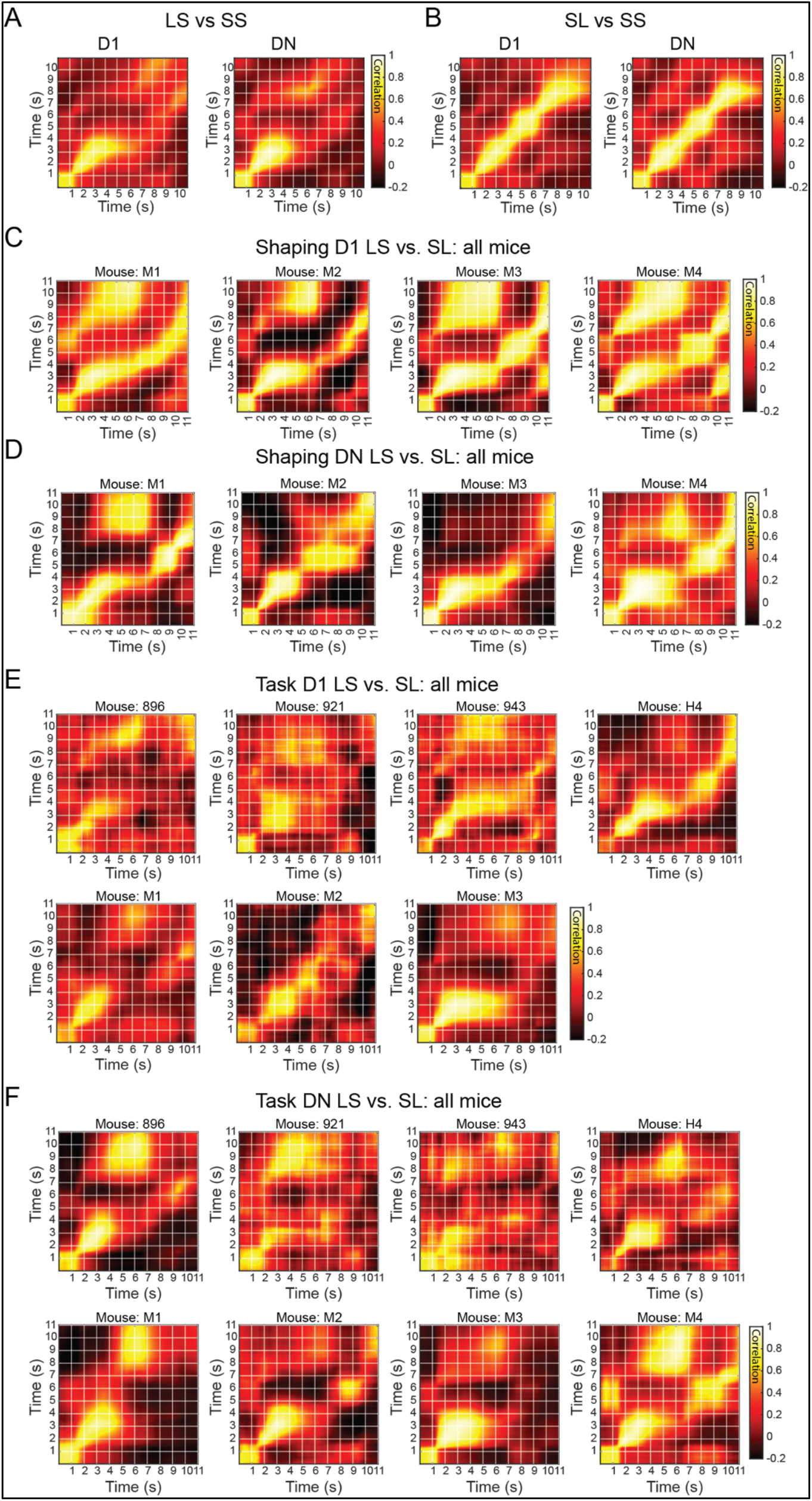
Additional cross-correlation matrixes. **A**. Population vector cross correlation matrices, shown for LS & SS trial types across training phases (from left to right-Task Dl, Task DN). **B**. Same as A but for SL & SS trial types. **C**. Population vector cross correlation matrices for each mouse, shown for LS & SL trial types on Shaping Dl. **D**. Same as C but for Shaping DN. **E**. Same as C but for Task Dl. **F**. Same as C but for Task DN.

## Methods

### Data collection

#### Subjects

All experiments were approved by and conducted in accordance with the University of Utah Animal Care and Use Committee. This study included 8 mice, aged 3-8 months. Mice were housed on a reverse 12-hour light/dark cycle in a room kept near 21.1°C with 25-45% humidity. All experiments were performed in the dark, during the dark phase of the light cycle. Mice were transgenic, expressing GCaMP6s for calcium imaging. Transgenic mice were created by crossing B6;DBA-Tg(tetO-GCaMP6s)2Niell/J and B6.Cg-Tg(Camk2a-tTA)1MmayDboJ (Jackson Laboratory) mice, and Camk2a-tTa;tetO-CGaMP6s double heterozygotes were used for experiments. Mice were water restricted prior to surgery and for the duration of experiments. Weight was carefully monitored to ensure mice maintained ≥80% of their baseline weight. Several mice (n=4) were imaged at all 4 timepoints. An additional 4 mice were only imaged on Task D1 and Task DN. One dataset was excluded from analysis, as the mouse was not engaged during Task D1.

#### CA1 implant

We followed previously established methods for CA1 implants (Dombeck et al 2010). Briefly, mice were anesthetized using 1-2% isoflurane. A craniotomy (∼2.8mm diameter) was made using a trephine drill over CA1 (x = 1.7, y = -2.3 relative to Bregma). Following the craniotomy, the dura was removed, then the cortex was slowly aspirated until the external capsule was exposed. The tissue was allowed to dry, then a cannula was inserted and fixed in place with Metabond. Cannulas were constructed by adhering a 2.5mm diameter round coverglass (Potomac) to a metal cannula (1.5mm height) with UV curable glue (Norland Optical Adhesive). Metabond was also used to attach a titanium headplate (10 x 40mm) to the skull, then a titanium ring (12.5mm inner diameter, 27mm outer) was attached to the headplate, centered over the cannula. Gaps between the skull, headplate, and objective ring were filled using Metabond.

#### Behavioral apparatus

All training was performed on one behavioral rig, positioned under a 2-photon microscope. Behavioral training was automated using an Arduino Uno, and data was collected at 1 kHz with a Picoscope Oscilloscope (Pico Technology v.6.13.2). Each session, mice were head-fixed over a treadmill (60cm circumference x 10cm wide) and were free but not required to run. At the start of each training session, a lick spout and odor delivery nozzle were positioned near the mouse. A capacitance sensor was used to detect licking (SparkFun Capacitive Touch), and solenoid valves were used to deliver water and odor when appropriate. Odors were delivered via a flow-dilution olfactometer, which combined a carrier (0.9 L/min) and odor stream (50mL/min) to deliver odor (2% isoamyl acetate in mineral oil). A solenoid valve controlled whether the odorized airflow was directed to the mouse or suctioned away by a vacuum (1.8 L/min).

#### tDNMS training

We followed previously established protocols for tDNMS training (Bigus et al. 2024). Briefly, training began with habituation, where mice were head-fixed and provided with water reward. Once mice reliably licked to take water (consuming ≥ 80% of drops in a series of 50 drops), they began the training process. Throughout training, each trial followed the same structure. Trial start was preceded by a flash of green light (3s prior to odor onset), then the first odor came on, followed by an interstimulus interval, followed by the second odor. A response window followed second odor offset. Trials were separated by an intertrial interval (16-24s). Mice were trained on one session per day, 5-7 days a week, typically for 100 trials during routine sessions, or for 65 minutes (∼130 trials) during imaging sessions.

Training began with shaping, where only nonmatched trials are presented. Trials were presented in a random order, with each trial type assigned a 50% probability of being chosen each trial. On Shaping D1 (Day 1 of shaping phase 1), a drop (6ul) of water was administered each trial 0.25s after second odor offset. Mice continued this training phase until they licked to take water in ≥ 80% of trials. Training then progressed to shaping phase 2, where mice had to lick in a 3s response window following second odor offset to earn reward. Water was administered after the first lick within the response window. If the mouse failed to earn reward, the next trial was automatically rewarded. Shaping phase 2 continued until mice successfully triggered reward in ≥ 20 consecutive trials. Upon reaching this criterion, mice advanced to shaping phase 3, where they had to withhold licking during the first odor and interstimulus interval (ISI), and lick during the response window to earn reward. As in shaping phase 2, each incorrect trial was followed by a trial with automatic reward delivery. Behavior was monitored until mice earned ≥ 20 consecutive rewards or reached 80% correct trials. Once mice met this benchmark, the next training session was used for imaging data collection and is referred to as Shaping DN. Following Shaping DN, mice advanced to the tDNMS task.

After shaping, mice began the tDNMS task, where matched trials were introduced. Imaging data were collected on the first day of tDNMS training (Task D1). Training continued until mice reached ≥75% correct trials, then a subsequent session was imaged and is referred to as Task DN. In the tDNMS task, matched trials are equally balanced with nonmatched trials. Each block of 4 trials contained 1 SL, 1 LS, and 2 SS trials presented in a random order within the block.

Correct SL and LS trials were rewarded with water; correct SS trials were not rewarded. Incorrect SL and LS trials were not punished; incorrect SS trials were punished with additional time (±12s) on the intertrial interval.

#### Two-photon imaging

A Neurolabware microscope with a 8-kHz resonant scanner was used to perform two-photon laser resonance scanning of CA1 neurons expressing GCaMP6s. Excitation was performed using a Ti:Sapphire laser (Discovery with TPC, Coherent) at 920nm. A 20x/0.45 NA air immersion objective (LUCPanFL, Olympus) was used, with an average power measured after the objective of 40-80mW. Data was collected via bidirectional scanning, with a frame rate of 30Hz. Scanbox software (v4.1) was used to acquire data, and imaging was also tracked on a Picoscope Oscilloscope sampling at 1kHz to align imaging and behavioral data. Behavioral data was later downsampled to align with imaging data.

#### Histology

Following behavioral experiments, perfusions were performed with 4% paraformaldehyde (PFA) in PBS (0.1M). After the perfusion, the brain was carefully removed and stored in 4% PFA (in 0.1M PBS) for 18-24 hours. Brains were then removed from PFA, rinsed 3x with PBS, and stored in PBS until sectioning. A vibrating microtome was used to create 100um coronal sections.

Sections were stored in PBS until staining. Prior to staining, sections were incubated in 0.1% Triton-X (in 0.1M PBS) for 15 minutes, then washed 3x with PBS. Immediately after PBS washes, sections were incubated with 29:1 solution of PBS (0.1M) and 435/455 blue fluorescent Nissl stain (Invitrogen) for 2-4 hours. Sections were then imaged using a VS200 Virtual Slide fluorescence microscope (Olympus).

### Data analysis

#### Time cell identification

Time cells were defined based on existing criteria (Heys & Dombeck 2018) with minor modifications. The activity of each cell was examined across all trials of a particular trial type (SL & LS in Shaping; SL, LS & SS in Task). We then calculated the mean dF/F of each cell in a particular context. Putative time fields were identified as continuous regions on this mean plot where activity was above a threshold of 50% of the difference between the peak dF/F value and the baseline value (average of the lowest 25% of dF/F values). To be classified as a time field, activity had to meet the following initial criteria: 1) fields must be ≥ 0.5s and ≤ 6s, 2) contain at lease one in-field value ≥ 6% dF/F 3) contain in-field activity in ≥33% of trials. If fields met initial criteria, mutual information was calculated as previously described (Bigus et al. 2024). For cells with multiple fields, activity in other fields was removed prior to mutual information analysis.

Briefly, to determine if mutual information of a putative time field was greater than expected by chance, dF/F values were shuffled circularly and mutual information was calculated for shufled data. Shufling was repeated 1000x, and the p-value was defined as the proportion of times that shufled mutual information was ≥ mutual information in the real data. Fields with a p value ≤ 0.01 were classified as time fields, and the following time field properties were recorded. First, the number of fields in each cell x context pair was noted. Cells could have multiple time fields in a context if each field met the above criteria. Time field start was defined as the first value on the mean dF/F plot that reached threshold (50% of the difference between the peak dF/F value and the lowest 25% of values). Similarly, field stop is when mean activity decreased below threshold. Field width is the time between field start and stop. Field peak refers to the time of the maximum mean dF/F value within the time field. Cells were classified as time cells if they had at least 1 significant time field in any context.

#### Time field properties

Time field width refers to the time between field start and stop, as mentioned above. Analysis of field width included data from all time fields that contained a field start and stop that was clearly within the boundaries of the segmented trial data. If the identified field start or stop was the first or last index, the field as excluded as we could not be sure when the field truly started or stopped. To analyze whether field width changes across the long odor, fields with activity in the SL and/or LS long odor were included. The time of each field’s peak was recorded relative to the long odor onset. All fields with peaks within each 1s bin of the long odor were included in analysis.

To determine when fields appeared in each session, we quantified the field onset trial number using established criteria (Dong et al. 2021). For each time field, we looked trial-by-trial (for all trials of a particular context) whether there was a significant transient within field boundaries (field start to stop). If so, we classified this as the putative onset trial and examined whether two of the next five trials of that trial type also had in-field activity. If so, the putative onset trial was classified as the field onset trial. If not, we repeated this analysis trial-by-trial until we identified field onset.

Previously described criteria were also used to classify whether each field shifted or was stable (Dong et al. 2021). For each time field, we first calculated the center of mass (COM) of each trial. Then, we performed linear regression using the matlab fitlm function. Fits with a p-value ≤ 0.05 were classified as shifting cells, and the shift was classified as negative or positive based upon the slope.

Finally, to analyze the number of fields, we determined which cell x context plots had significant time fields. Only these data were considered. We then quantified whether those cell x context plots had one or more time fields, defined using the above criteria.

#### Population activity

Cross correlation matrices were used to compare time cell activity between trial types. Activity of all time cells from all mice was included in analysis. First, the average activity of each time cell was found across all trials of a context. These values were used to create population vectors, which were then correlated with Pearson’s correlation coefficient. To quantify the correlation between trial types in key periods, we took the average of all correlation values within the time bins that defined the area of interest.

PCA plots were generated by performing principal component analysis on the trial averaged activity of time cells. Average activity of each time cell in each trial type was included to find variability between trial types. Data were projected on the first two principal components. Data were binned in 1s bins then connected with lines for visualization. While all other analysis was performed on dF/F data, PCA was performed with deconvolved data.

#### Variance in time or space

For each trial, the peak dF/F location was measured, either relative to distance travelled or time elapsed (beginning 1s before odor 1 onset). The variability of peak locations (in distance or in time) across trials was assessed using the coefficient of variation (ratio of the standard deviation to the mean). Values were compared in the distance and time dimensions with a Student’s Paired t-test.

#### Correlations between trial types

To examine whether time cells display similar activity across pairs of contexts, all time cells with fields in the compared trial types were selected. Pearson’s correlation coefficient was used to determine the extent of similarity in mean activity between contexts.

#### Time of divergence

To further characterize the extent of similarity or differences in time cell activity between contexts, we aimed to identify the time of divergence, or the time when population level time cell activity became significantly different across contexts. First, cells with time fields in the selected contexts were identified. Population vectors were created for each context, and the difference in dF/F was determined for each time bin. This real difference in data was compared to a shufle distribution, generated by randomly assigning trial types and calculating the dF/F difference 1,000 times. The time of divergence is the time when the real difference surpasses the top 0.1% of the shufled distribution.

#### Remapping

For cells with multiple fields, only the field with the highest amplitude peak was used for analysis. All cells with time fields in SL or LS trials were included in analysis. It was first determined if there was a field in both contexts. If not, the time cell was classified as losing a field (field disappeared) in one context. If fields were present in each context, it was determined where the field was stable or remapped. Fields were classified as stable if the peak in one context was within 0.5s in either direction (before or after) of the peak in the other context. If this criteria was not met, the field was classified as remapping. Fields remapped to the same phase if the peaks for each context were in the same task phase (ex: each peak was in odor 1; each peak was in the ISI; each peak was in odor 2), or different phases (for instance, if one peak was odor 1 and the other was odor 2).

#### Behavior

Correct nonmatch trials were defined as those where mice withheld licking in the first odor and interstimulus interval (ISI) and licked in the response window (3s) following second odor offset to successfully trigger reward. Correct match trials were defined as those where mice withheld licking during the trial period and response window. To further characterize licking behavior, we examined the percent of trials in which mice licked following first odor offset, specifically in the 3s ISI. Additionally, we examined predictive licking, defined by licking within the last 0.5s before second odor offset. Velocity was also examined. Due to different running speeds of different mice, velocity was normalized to the maximum for each mouse within the session.

### Statistics

Sample sizes were based on previous studies (Bigus et al. 2024; Heys et al. 2018) and not predetermined with statistical methods. Statistical tests were used when applicable and included Kruskal-Wallis tests (with Dunn-Bonferroni post-hoc testing), linear mixed effects models, and paired t-tests. Linear mixed effects models were used to compare values from individual mice across days, since sample sizes varied as we added mice on Task D1. For linear mixed effects models, the matlab function fitlme was used, followed by pairwise comparisons (with Bonferroni correction).

